# *Salmonella* manipulates the host to drive pathogenicity via induction of interleukin 1β

**DOI:** 10.1101/2023.06.14.544934

**Authors:** Mor Zigdon, Jasmin Sawaed, Lilach Zelik, Dana Binyamin, Shira Ben-Simon, Nofar Asulin, Rachel Levin, Sonia Modilevsky, Maria Naama, Shahar Telpaz, Elad Rubin, Aya Awad, Wisal Sawaed, Sarina Harshuk-Shabso, Meital Nuriel-Ohayon, Michal Werbner, Omry Koren, Sebastian E Winter, Ron N Apte, Elena Voronov, Shai Bel

## Abstract

Acute gastrointestinal infection with intracellular pathogens like *Salmonella* Typhimurium triggers the inflammasome and the release of the proinflammatory cytokine interleukin 1β (IL-1β). However, the role of IL-1β in intestinal defense against *Salmonella* remains unclear. Here, we show that IL-1β production is detrimental during *Salmonella* infection. Mice lacking IL-1β (*IL-1β* ^-/-^) failed to recruit neutrophils to the gut during infection, which reduced tissue damage and prevented depletion of short-chain fatty acid-producing commensals. Changes in epithelial cell metabolism that typically support pathogen expansion, such as switching energy production from fatty acid oxidation to fermentation, were absent in infected *IL-1β*^-/-^ mice which inhibited *Salmonella* expansion. Additionally, we found that IL-1β induces expression of complement anaphylatoxins and suppresses the complement-inactivator Carboxypeptidase N (CPN1). Disrupting this process via IL-1β loss completely prevented mortality in *Salmonella*-infected *IL-1β*^-/-^ mice and led to chronic infection. Thus, *Salmonella* exploits IL-1β signaling to outcompete commensal microbes and establish gut colonization. Moreover, our findings identify the intersection of IL-1β signaling and the complement system as key host factors involved in controlling mortality during invasive Salmonellosis.

## Introduction

The cytokine interleukin 1β (IL-1β) is a proinflammatory alarmin which is released mainly from activated myeloid cells during acute and chronic inflammation. To facilitate its rapid secretion, IL-1β is stored in the cell as a propeptide which is cleaved to its mature form by caspase 1 following activation of the inflammasome by invasive bacteria. Once secreted, IL-1β affects multiple cell types via binding to the IL-1 receptor(*1*). Notably, IL-1β secretion affects endothelial cell permeability to allow massive infiltration of neutrophils from the periphery into the infected tissue(*2*). While these activated neutrophils contribute to bacterial killing, they also have a damaging effect on host tissues.

The foodborne pathogen *Salmonella* enterica serovar Typhimurium (hereafter *Salmonella*) is a common cause of acute gastrointestinal inflammation, caused by consumption of contaminated food. While this infection is usually self-limiting in healthy humans, it can lead to life-threatening bacteremia in immune-compromised individuals. In mice carrying a mutated *Nramp1* gene, such as all mice on a C57BL/6 background, oral infection leads to acute colitis and systemic dissemination of *Salmonella*, ultimately resulting in death within several days(*3*). The *Nramp1* gene encodes an ion channel expressed in macrophages and it is thought that loss of this gene renders phagocytic cells unable to control growth of intracellular bacteria(*4*). Yet the mechanism leading to mortality caused by *Salmonella* infection in mice is not clear.

Studies in mice have revealed that *Salmonella* can exploit the host’s innate immune responses to allow its own expansion. Mice lacking toll-like receptors (TLRs) 2, 4 and 9 are more resistant to *Salmonella* infection than mice lacking either TLRs 2 and 4 or 4 and 9 alone, indicating that bacterial sensing by the host can be detrimental(*5*). In another example, mice lacking the cytokine IL-22 were found to limit *Salmonella* luminal growth better than wild type mice (WT), as secretion of lipocalin-2 and calprotectin under the control of IL-22 suppressed commensal microbes which directly compete with *Salmonella*(*6*). Thus, *Salmonella* has evolved to manipulate the host’s defense mechanisms for its own advantage(*7*). Here, we set out to determine whether host production of IL-1β is protective during *Salmonella* infection and discovered that the pathogen exploits this cytokine to drive pathogenicity.

## Results

### Loss of IL-1β in mice reduces Salmonella burden and prevents mortality during oral infection

Previous studies have shown that IL-1β has a protective role during chemical-induced colitis, by driving epithelial repair following injury(*8*). To determine whether IL-1β is required for host defense during infection-induced colitis we challenged streptomycin-treated WT and *IL-1β*^-/-^ mice (on a C57BL/6 background) with a single oral dose of *Salmonella* (Figure 1A).

**Figure 1.**
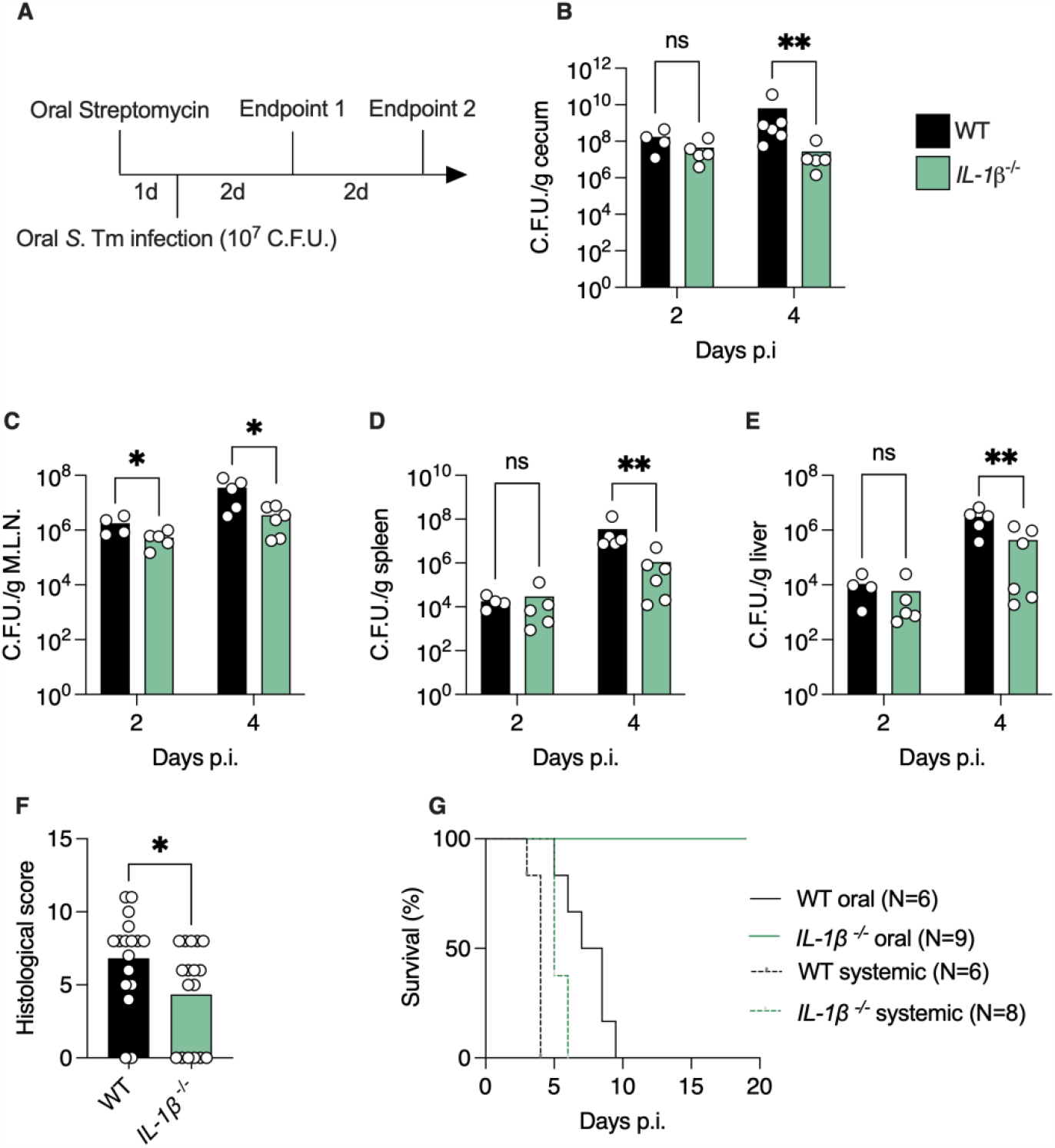
Loss of IL-1β dampens *Salmonella* growth *in vivo* and prevents mortality from oral infection. (**A**) Scheme depicting experimental design. (**B**-**E**) *Salmonella* C.F.U. in cecum (**B**), M.L.N. (**C**), spleen (**D**) and liver (**E**) of mice infected as in (**A**). (**F**) Histological damage in colonic tissue 2 d.p.i. (**G**) Survival percentage of mice infected orally or intravenously (systemic infection) with *Salmonella*. (**B-F**) Each dot represents a mouse. \**P*<0.05; \*\**P*<0.01; Mann-Whitney test. C.F.U., colony-forming units; p.i., post-infection.

Pretreatment with streptomycin is crucial to drive robust gastrointestinal inflammation(*9*). We found that *Salmonella* levels were reduced 100-fold in the gut lumen of *IL-1β*^-/-^ mice 4 days post-infection (d.p.i.) compared to WT mice (Figure 1B). *IL-1β*^-/-^ mice also harbored 10-fold less *Salmonella* in gut-draining mesenteric lymph nodes (M.L.N.), spleen, and liver compared to WT (Figure 1C-E). This reduction in bacterial loads in *IL-1β*^-/-^ mice was accompanied by a marked reduction in histological colonic tissue damage (Figure 1F). Thus, loss of IL-1β limits *Salmonella* expansion in the gut lumen and systemic sites.

We then tested whether loss of *IL-1β* affects infection-induced mortality. Remarkably, we found that *IL-1β*^-/-^ mice were completely resistant to infection-induced mortality, while WT mice all succumbed to the infection (Figure 1G). To determine whether this resistance to infection is dependent on mode of infection we infected mice intravenously with *Salmonella*.

Previous reports have established that mice carrying the resistance *Nramp1* allele do not succumb to low-dose intravenous infection with *Salmonella*, while mice on a C57BL/6 background die within a week(*4*). We found that *IL-1β*^-/-^ mice infected intravenously with a low dose of *Salmonella* die from the infection in the same manner as WT mice (Figure 1G). These results indicate that IL-1β plays a role in mortality during oral *Salmonella* infection.

### IL-1β^-/-^ mice fail to recruit neutrophil to the gut during Salmonella infection

We next wanted to determine how IL-1β production drives *Salmonella* expansion in the gut lumen. Mice carrying the mutant *Nramp1* allele, such as C57BL/6 mice, are thought to fail in controlling *Salmonella* expansion because of defective macrophage function(*3*). Thus, we hypothesized that IL-1β impairs intracellular killing of *Salmonella* by macrophages. To test this, we extracted peritoneal macrophages from WT and *IL-1β*^-/-^ mice and tested their ability to kill intracellular *Salmonella in vitro*. Contrary to our hypothesis, we found that macrophages from *IL-1β*^-/-^ mice were less efficient at killing intracellular *Salmonella* (Figure S1A). This result indicates that loss of IL-1β impairs *Salmonella* growth in a manner which is not dependent on intracellular killing by macrophages.

To understand how IL-1β deficiency affects the colonic tissue during *Salmonella* infection we performed bulk RNA sequencing followed by pathway analysis of differentially expressed genes. We found that genes which were downregulated in *IL-1β*^-/-^ mice are involved in neutrophil function and recruitment to the colon (Figure 2A). Indeed, key chemokines which attract neutrophil and monocytes to the colon, along with neutrophil-specific transcripts such as lipocalin-2 and calprotectin, were expressed at lower levels in *IL-1β*^-/-^ mice (Figure 2B). We validated these findings by counting myeloperoxidase (MPO)- and neutrophil elastase (NE)-positive cells in colonic tissue sections from infected mice. Accordingly, *IL-1β*^-/-^ mice had on average 4-fold fewer neutrophils in their colons (Figure 2C and D). We then tested whether loss of IL-1β leads to reduced neutrophils levels during *Salmonella* infection because of neutrophil development defects, or failure to recruit neutrophils to the colonic tissue. By analyzing circulating blood from naïve and infected mice, we found that loss of IL-1β does not affect circulating neutrophil levels during steady state (Figure S2A), but rather prevents neutrophil recruitment during infection (Figure 2E). Thus, the resistance of *IL-1β*^-/-^ mice to *Salmonella* is not due to an elevated antimicrobial response as they fail to draw neutrophils to the site of infection.

**Figure 2:**
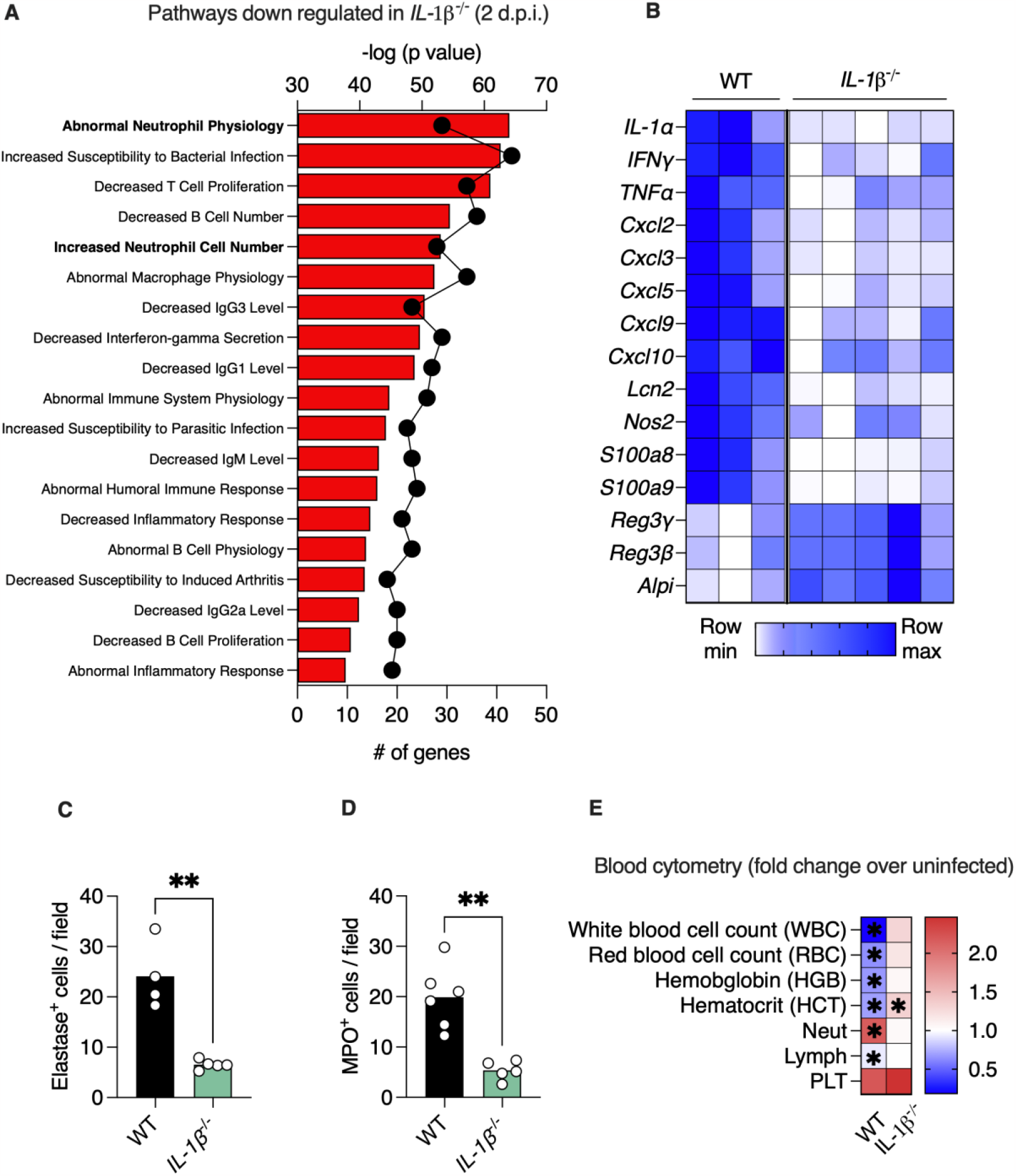
Loss of IL-1β impairs neutrophil recruitment to the gut during *Salmonella* infection. (**A**) Pathway analysis of transcripts which are downregulated in colonic tissue of *Salmonella*-infected *IL-1β*^-/-^ according to GO biological function. Bars represent -log (*P* value) and dots represent number of genes in pathway. (**B**) Heatmap depicting differentially expressed innate immune genes with a *P*<0.05. Each column represents a mouse and each row a gene. (**C-D**) Numbers of neutrophil elastase-positive (**C**) and myeloperoxidase-positive (**D**) cells in colonic section of *Salmonella*-infected mice. Each dot represents a mouse. (**E**) Heatmap depicting fold-increase (infected/naïve) in levels of indicated cell types in whole blood. \**P*<0.05; \*\**P*<0.01; Student’s t-test. d.p.i., days post-infection; Neut, neutrophils; Lymph, lymphocytes; PLT, platelets.

### IL-1β production leads to collapse of gut short-chain fatty acid-producing Clostridia during Salmonella infection

Commensal gut microbes provide colonization resistance by directly inhibiting pathogen growth and by affecting the host(*10*). We next tested whether changes in the gut microbiota of *IL-1β*^-/-^ mice can explain their resistance to *Salmonella* infection. We reasoned that lack of neutrophil recruitment to the gut would dampen the harmful damage of infection-induced inflammation on the microbiota(*10*). 16S rRNA microbiota analysis of feces from infected WT and *IL-1β*^-/-^ mice revealed that the latter contained a more diverse microbiota after *Salmonella* infection (Figure 3A). Jaccard analysis confirmed that the gut microbial composition of infected *IL-1β*^-/-^ mice was distinct from that of WT mice (Figure 3B). Analysis of composition of microbiomes (ANCOM) showed that bacteria from the Clostridia class were enriched in the gut of infected *IL-1β*^-/-^ mice while these microbes were virtually absent from the gut of infected WT mice (Figure 3C). Previous reports have shown that *Salmonella* infection leads to depletion of short-chain fatty acid (SCFA)-producing Clostridia microbes(*11*). We found that in naïve WT and *IL-1β*^-/-^ mice the levels of Clostridia bacteria were similar. However, *Salmonella* infection dramatically reduced the levels of Clostridia bacteria in infected WT mice while not affecting these microbes in *IL-1β*^-/-^ mice (Figure 3D). Thus, *Salmonella* infection does not lead to depletion of SCFA-producing Clostridia bacteria in *IL-1β*^-/-^ mice.

**Figure 3:**
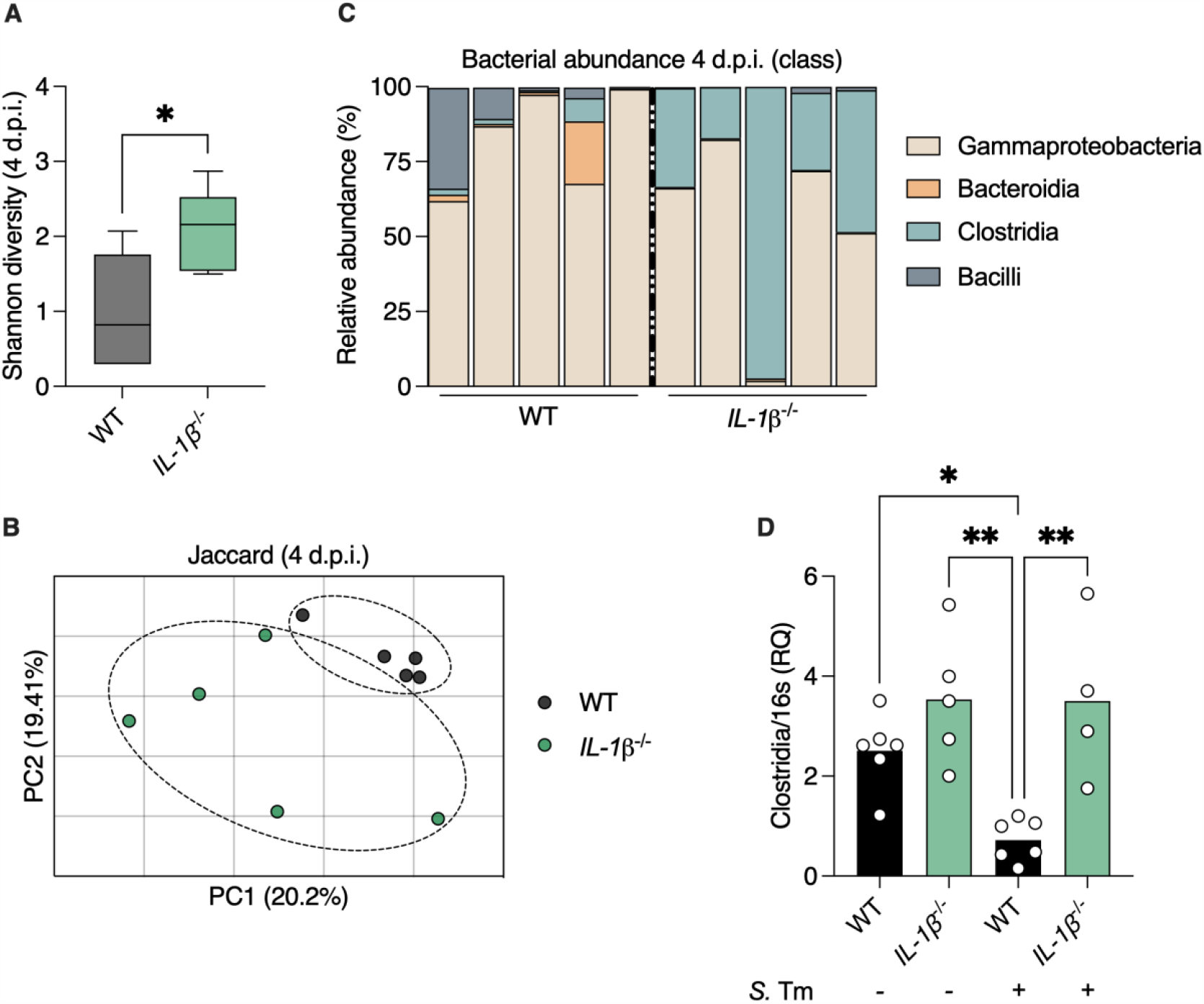
*Salmonella* infection does not deplete SCFA-producing Clostridia from the gut of *IL-1β* ^*-/-*^ mice. 16 S rRNA sequencing was performed to characterize gut microbiota composition. (**A**) Shannon index representing microbial diversity in gut microbiota of *Salmonella*-infected mice. (**B**) PCoA of fecal microbiota based on Jaccard similarity coefficient. Each dot represents a mouse. (**C**) Relative taxonomic composition at the Class levels. Each column represents a mouse. (**D**) qPCR analysis of levels of the class Clostridia in naïve or infected mice as indicated. Each dot represents a mouse. \**P*<0.05; \*\**P*<0.01; (**A**) Student’s t-test, (**D**) One-way ANOVA. d.p.i., days post-infection; *S*. Tm, *Salmonella* typhimurium.

### Preservation of SCFA-producing bacteria in IL-1β^-/-^ mice inhibits Salmonella expansion by maintaining beta oxidation in colonocytes and hypoxic condition in the gut

Next, we wanted to determine how this different microbiota in *IL-1β*^-/-^ mice affects the colonic tissue. We found that genes which were upregulated in the colon of infected *IL-1β*^-/-^ mice were enriched in the metabolism pathway (Figure 4A). Specifically, we found that genes encoding proteins that participate in energy production via fatty acid beta oxidation were upregulated in infected *IL-1β*^-/-^ mice as compared to infected WT mice (Figure 4B).

**Figure 4:**
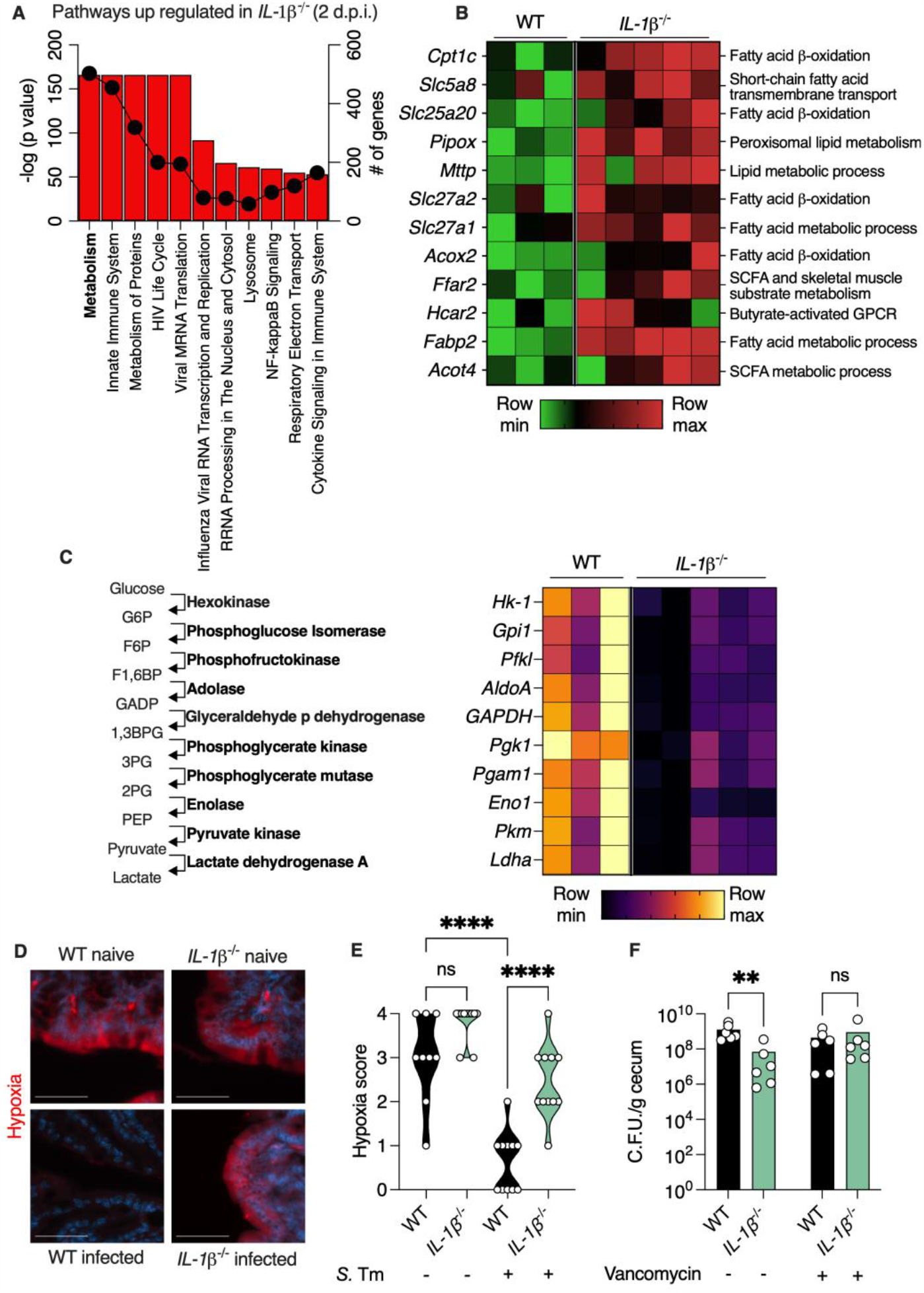
Preservation of fatty acid beta-oxidation in *IL-1β* ^*-/-*^ mice inhibits *Salmonella* growth *in vivo*. (**A**) Pathway analysis of transcripts which are upregulated in colonic tissue of *Salmonella*-infected *IL-1β*^-/-^ according to GO biological function. Bars represent -log (*P* value) and dots represent number of genes in pathway. (**B**) Heatmap depicting differentially expressed genes involved in fatty acid oxidation with a *P*<0.05. Each column represents a mouse and each row a gene. (**C**) Heatmap depicting differentially expressed genes in the glycosylation pathway with a *P*<0.05. The enzymatic activity of each gene in the glycosylation pathway is presented on the left. Each column represents a mouse and each row a gene. (**D**) Immunofluorescence microscopy of colonic section from mice. Red staining shows pimonidazole which indicates hypoxia levels. Nuclei were stained with DAPI. Scale bar, 50μm. (**E**) Quantification of red signal in (**D**). Each dot represents a mouse. (**F**) *Salmonella* C.F.U. in cecum of infected mice 4 d.p.i. treated as indicated. Each dot represents a mouse. \*\**P*<0.01, \*\*\*\**P*<0.0001; (**E**) One-way ANOVA; (**F**) Mann-Whitney test. d.p.i., days post-infection; *S*. Tm, *Salmonella* typhimurium.

SCFAs produced by commensal Clostridia bacteria serve as the preferred energy source for colonocytes. These epithelial cells produce ATP via beta oxidation of SCFA, which renders the colonic lumen hypoxic as this process utilizes oxygen. *Salmonella* infection has been shown to deplete SCFA-producing bacteria from the colon, thus forcing colonocytes to produce energy via glycosylation(*11*). As this process does not require oxygen, *Salmonella* infection effectively leads to elevated oxygen levels in the colonic lumen. Oxygen in the lumen is toxic to the obligatory anaerobic commensal microbiota, which is beneficial for *Salmonella* as it disrupts the colonization resistance provided by these commensals(*10*). Indeed, we found that all the genes in the glycosylation pathway were elevated in infected WT compared to *IL-1β*^-/-^ mice (Figure 4C). Accordingly, we found that the colonocytes in *IL-1β*^-/-^ mice remained hypoxic during *Salmonella* infection while colons of WT mice became rich in oxygen (Figure 4D and E).

These observations led us to hypothesize that failure to recruit neutrophils in *IL-1β*^-/-^ mice preserves hypoxic conditions in the colon by sparing SCFA-producing Clostridia bacteria. This in turn can inhibit *Salmonella* growth by providing direct competition and by reducing oxygen levels which are important for *Salmonella* growth(*10*). To test our hypothesis, we depleted Clostridia bacteria from the colon of WT and *IL-1β*^-/-^ mice by vancomycin treatment and then infected these mice with *Salmonella*. Indeed, we found that *Salmonella* growth in the intestine of *IL-1β*^-/-^ mice was indistinguishable from WT mice after vancomycin treatment (Figure 4F). Thus, preservation of SCFA-producing bacteria in *IL-1β*^-/-^ mice maintains energy production via beta oxidation and hypoxic condition in the gut which in turn inhibit *Salmonella* growth.

### IL-1β drives mortality in Salmonella-infected mice by suppressing the anaphylatoxin-inactivator Carboxypeptidase N

Next, we wanted to determine whether preserving SCFA-producing Clostridia is the mechanism which prevents mortality in *Salmonella*-infected *IL-1β*^-/-^ mice. However, we found that vancomycin-treated *IL-1β*^-/-^ mice still survived after oral *Salmonella* infection (Figure 5A). We then hypothesized that the reason for resistance to *Salmonella* infection in *IL-1β*^-/-^ mice is that they can clear the infection. Yet we found that even 17 d.p.i. these *IL-1β*^-/-^ mice were still colonized by *Salmonella* (Figure 5B), and that the pathogen was still fully virulent as transmission to a WT host was still lethal (Figure 5C).

**Figure 5:**
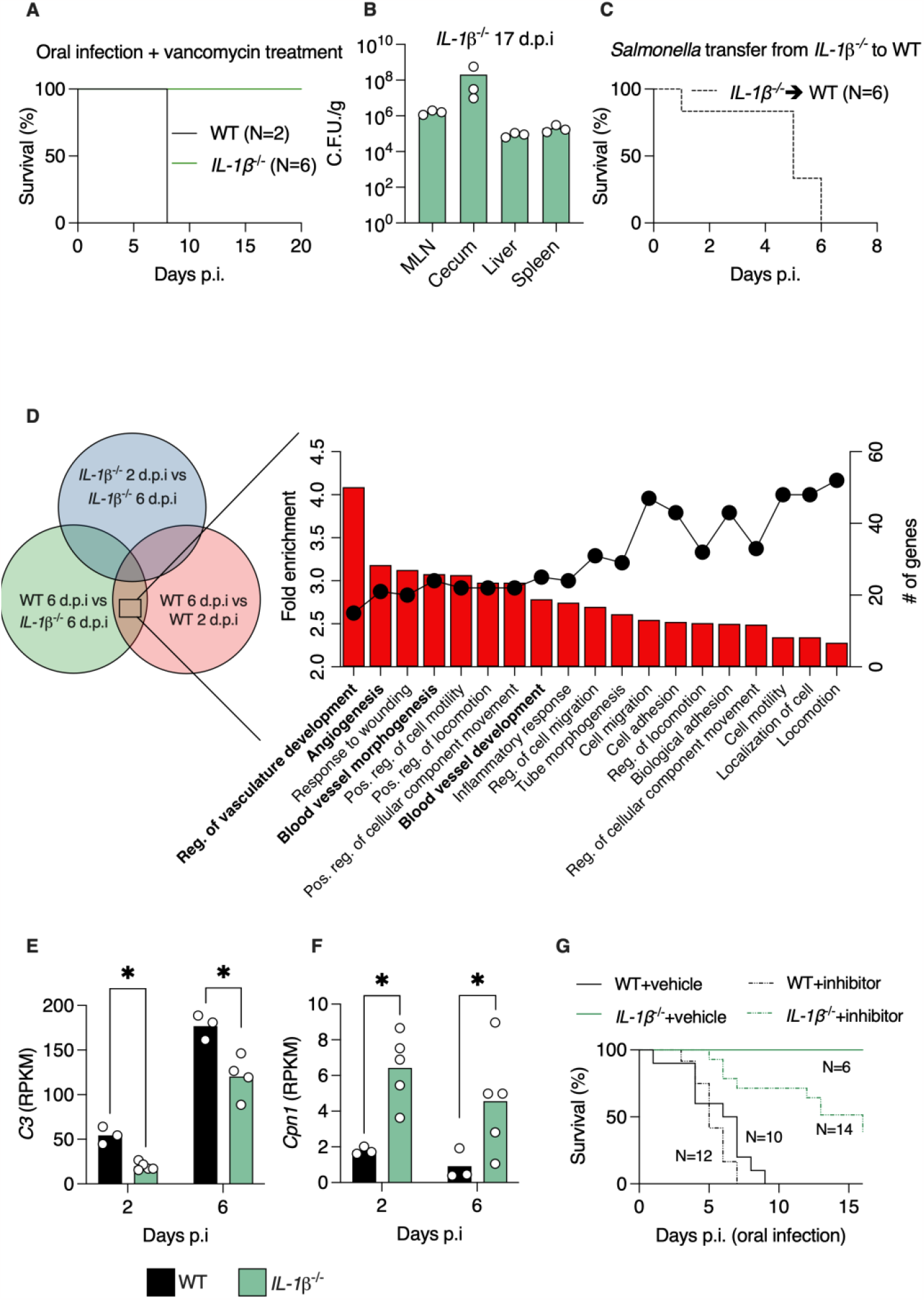
IL-1β promotes expression of anaphylatoxin and inhibits expression of carboxypeptidase N which drives mortality in *Salmonella* infected mice. (**A**) Survival of vancomycin-treated mice infected orally. (**B**) *Salmonella* C.F.U. in the indicated organs of *IL-1β*^-/-^ mice 17 d.p.i. Each dot represents a mouse. (**C**) Survival of WT mice infected with *Salmonella* isolated from *IL-1β*^-/-^ mice 21 d.p.i. (**D**) Pathway analysis of transcripts which are differently expressed as indicated in Venn diagram in colonic tissue of *Salmonella*-infected mice according to GO biological function. Bars represent fold enrichment and dots represent number of genes in pathway. (**E-F**) Normalized reads of the indicated genes from colons of mice based on RNA sequencing. Each dot represents a mouse. (**G**) Survival of *Salmonella*-infected mice treated with vehicle or CPN1 inhibitor. \**P*<0.05; (**E-F**) Student’s t-test. d.p.i., days post-infection; CPN, carboxypeptidase N.

**Figure 6:**
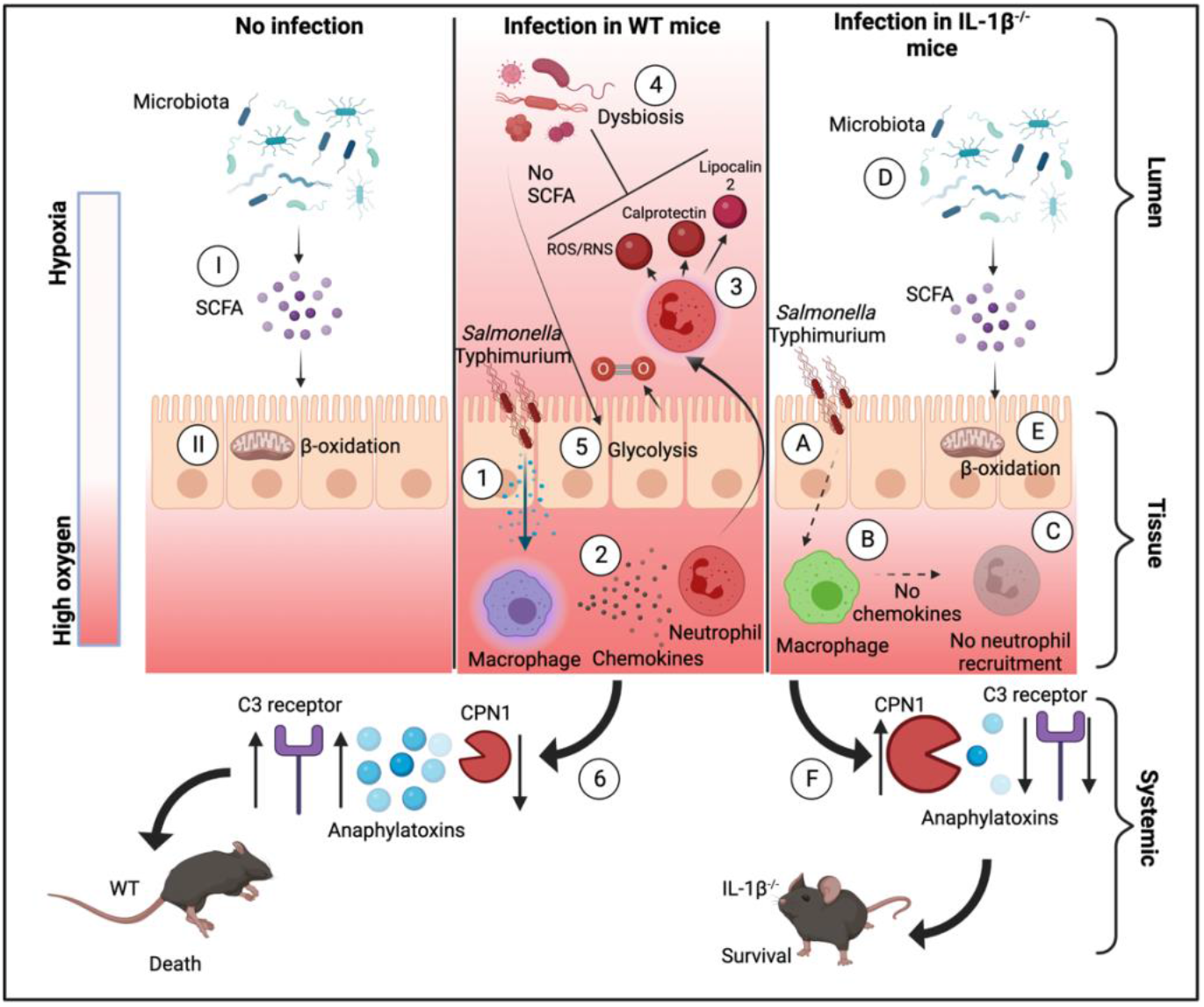
IL-1β is a key axis in host manipulation by *Salmonella*. (**I**) In naïve mice, commensal microbes produce SCFA. (**II**) These SCFA are used to produce energy via β-oxidation by colonocytes. As this process uses oxygen, the gut lumen remains hypoxic. (**1**) *Salmonella* infection in WT mice leads to IL-1β secretion and (**2**) production of chemokines that attract neutrophil to the gut lumen. (**3**) These neutrophil produce ROS and NOS and secrete metal chelators which (**4**) kills commensal microbes, thus reducing SCFA levels. This forces colonocytes to produce energy via fermentation which does not uses oxygen, thus allowing oxygen to seep into the gut lumen, further damaging the microbiota. (6) Release of IL-1β following infection leads to elevated levels of anaphylatoxins and suppressed levels of CPN1, leading to mortality. (**A**) In mice lacking IL-1β, *Salmonella* infection (**B**) does not lead to production of chemokines and (**C**) does not attract neutrophils to the gut lumen. (**D**) These preserve SCFA-producing commensals and (**E**) allow colonocytes to produce energy via β-oxidation, which preserves luminal hypoxia which impedes *Salmonella* expansion. (**F**) Lack of IL-1β reduces anaphylatoxin levels and maintains high levels of CPN1 which prevents mortality.

To understand why *IL-1β*^-/-^ mice do not die from *Salmonella* infection we performed RNA sequencing on mice 6 d.p.i., as this is the timepoint when WT mice are moribund. We reasoned that genes which are differently expressed in WT mice 6 d.p.i. compared to WT mice 2 d.p.i., and to *IL-1β*^-/-^ mice 6 d.p.i., are related to the process that leads to mortality. We excluded genes that are differently expressed in *IL-1β*^-/-^ mice 6 d.p.i. versus 2 d.p.i. as these mice do not die from the infection, thus these genes are not related with mortality (Figure 5D). Pathway analysis revealed that WT mice express genes that are related to vasculature morphology 6 d.p.i. (Figure 5D). We found this interesting, as septic shock which occurs during systemic infection by pathogens is characterized by vasculature permeability, dilation, and a severe hypotensive response(*12, 13*). These harmful changes to the vasculature are mediates by complement proteins, also known as anaphylatoxins, which can affect vasculature permeability at sub-nanomole concentrations(*12*). We found that the complement protein C3, and the complement receptor C3ar1 were highly induced in WT mice at 6 d.p.i. (Figure 5E and Figure S3A). While these genes were also induced in *IL-1β*^-/-^ mice, their levels at 2 and 6 d.p.i. were still significantly lower than in WT mice (Figure 5E and Figure S2A). Thus, IL-1β drives expression of anaphylatoxin proteins and receptors during *Salmonella* infection.

Under normal conditions, the damaging effects of anaphylatoxins are balanced by carboxypeptidase N (CPN1), which inactivates anaphylatoxins by peptide cleavage within seconds at physiological concentrations(*14, 15*). We found that *IL-1β*^-/-^ mice express 4-fold higher levels of *Cpn1* after *Salmonella* infection, compared with WT mice (Figure 5F). Indeed, a previous report has shown that IL-1 receptor signaling inhibits *Cpn1* expression(*16*). These results imply that low expression and inactivation of the complement system in *IL-1β*^-/-^ mice preserves their viability during infection. To test this, we treated mice with a CPN1 inhibitor. We found that inhibiting CPN1 in uninfected mice does not affect their viability (Figure S4A). In infected WT mice, administration of CPN1 inhibitor did not affect survival (Figure 5G). However, infected *IL-1β*^-/-^ mice treated with CPN1 inhibitor suffered from over 60% mortality, while vehicle-treated *IL-1β*^-/-^ mice were completely resistant to infection-induced mortality (Figure 5G). Thus, loss of IL-1β protects mice from *Salmonella*-induced mortality by removing suppression from CPN1 expression.

## Discussion

*Salmonella* has evolved to be a master manipulator of host response. By driving acute inflammation in the gut, *Salmonella* facilitates the killing of its microbial competitors via the host. Indeed, recent research has shown that, under experimental settings, an immune-compromised host (through loss of TLR signaling, IL-22 production or STAT2 signaling) is better equipped to contain *Salmonella* infection(*5, 6, 17*). Here, we show that IL-1β is a central component of host-manipulation by *Salmonella*. Using IL-1β signaling to facilitate its colonization is an effective mechanism deployed by *Salmonella*, as multiple microbial detection apparatuses (TLRs and inflammasomes) converge on processing and release of IL-1β(*18*). This capability of different pathways to drive IL-1β secretion can also explain why mice deficient in individual components of the inflammasome have not shown the same resistance to *Salmonella* infection as we show here in *IL-1β*^-/-^ mice(*19*). Along these lines, a recent report has shown that only simultaneous ablation of all the pathways which drive NLRC4 inflammasome-dependent cell death can protect mice from fatal infection(*20*). Thus, it is possible that *Salmonella* has evolved to depend on IL-1β for host colonization, as infection consistently results in IL-1β secretion, even if not all pathogen-detection mechanisms are triggered.

Many studies in the fields of immunology and pathogenesis rely on mouse mortality as a measurable readout for virulence. In the current study, we found that *IL-1β*^-/-^ mice do not succumb to *Salmonella* infection. To help us identify the mechanism leading to this resistance, we conducted a literature search to understand why this infection is lethal in C57BL/6 WT mice. Yet we were surprised that our literature search on the mechanisms that drive mortality during non-typhoidal Salmonellosis did not result in definitive explanations. Death from non-typhoidal Salmonellosis in mice carrying the susceptibility allele in *Nramp1* is attributed to an inability to control intracellular bacteria by macrophages(*4*). Yet we could not find definitive evidence to support this notion. Another study has found that mortality from intravenous infection is dependent on bacterial lipid A and is characterized by robust IL-1β secretion(*21*). Yet the downstream target of IL-1β which leads to mortality was not known. Our discovery that IL-1β dramatically upregulates the expression of complement proteins and downregulates the expression complement-inactivator *in vivo* provides a possible mechanism for *Salmonella*-driven mortality. This mode of regulation can thus maximize the antimicrobial activity of the complement system as it drives complement expression and inhibits its degradation simultaneously.

In humans, a global systematic review and meta-analysis found that the most common reason for non-typhoidal *Salmonella* disease-associated complications and fatalities was septicaemia(*22*). The definition of septicaemia is “an infection that occurs when bacteria enter the bloodstream and spread”, according to the Cleveland Clinic(*23*). As in mice, this explanation does not provide a testable mechanism. Studies in non-human primates have shown that inhibition of complement proteins can prevent mortality from intravenous bacterial infection(*24*). Complement proteins are known to elevate vascular permeability, cause smooth muscle contraction and drive cardiomyopathy, all of which lead to fatal outcome(*25*). Thus, control of complement expression needs to be linked to acute immune activation, while also being tightly regulated to prevent excessive damage. Our finding that IL-1β drives expression of complement anaphylatoxins, and suppresses expression of complement-inactivating CPN1, provides an example where immune regulation of the complement system can spiral out of control, leading to death. This also provides a possible mechanism for non-typhoidal *Salmonella*-induced mortality. Whether this is the mechanism which results in fatalities from septicaemia in humans will need to be tested.

## Acknowledgments

This study was performed in memory of Ron N Apte.

## Funding

The Azrieli Foundation Early Career Faculty Fellowship (SB)

Israeli Science Foundation (ISF) grants 925/19 and 1851/19 (SB)

European Research Council (ERC)

Starting Grant GCMech 101039927 (SB)

U.S-Israel Binational Science Foundation 2021025 (SEW and SB)

## Author contributions

Conceptualization: MZ, MW, SEW, RNA, EV, and SB.

Investigation: MZ, JS, LZ, DB, SBS, NA, RL, SM, MN, ST, ER, AA, MNO, and MW.

Funding acquisition: SEW, RNA, EV, and SB. Supervision: OK, SEW, RNA, EV, and SB. Writing – original draft: SEW, EV, and SB. Writing – review & editing: SEW, EV, and SB.

## Materials and Methods

### Mice

C57BL/6 wild-type mice and *IL-1β*^-/-^ mice (backcrossed to C57BL/6 for over 10 generations) were separately bred and maintained in the conventional barrier at the Azrieli Faculty of Medicine, Bar-Ilan University, Israel. 8–14-week-old mice were used for all experiments. All experiments were performed using protocols approved by the Institutional Animal Care and Use Committees (IACUC) of the Bar-Ilan University.

### Salmonella infection

Mice were treated with 20mg streptomycin via gavage 24 hours before oral infection with 10^7^ C.F.U. *Salmonella* enterica serovar typhimurium (SL1334) or intravenous infection with 10^5^ C.F.U. *Salmonella*. For vancomycin treatment model, mice were treated with 300mg of streptomycin and 150mg of vancomycin in drinking water in a volume of 300ml throughout the course of the infection. At the designated timepoints mice were euthanized and cecal content, liver, spleen, and mesenteric lymph nodes were removed, weighed, homogenized in PBS, and plated on LB agar plates containing streptomycin following serial dilutions. For survival assays, mice that were unresponsive to touch, cold or could not reach their food were euthanized.

### Gentamycin protection assay

Mice were treated with 1ml of thioglycolic acid via intraperitoneal injection and euthanized after 5 days, after which 5ml of PBS was injected to the peritoneal cavity and mixed gently. The PBS containing cells was then drawn out and stored on ice. The solution was centrifuged for 10 minutes at 400g, supernatant discarded, and cell pellet was resuspended with 1ml DMEM with serum and filtered in 40um filter. Filtered cells were counted and 10^6^ cells were plated on cell culture plate overnight at 37^0^C. Wells were washed 3 time with PBS (37^0^C) and incubated with *Salmonella* at a multiplicity of infection (MOI) =3 for 90 minutes. Infected cells were washed 3 time with PBS (37^0^C) and incubated with 200mg/ml gentamicin for 90 minutes and then washed with PBS. Lysis was then performed with 1% of Triton X-100 for 15 minutes. The lysate was diluted and seeded on LB agar plates containing streptomycin for overnight incubation.

### RNA sequencing and analysis

RNA from frozen colonic tissues was extracted using Qiagen RNeasy Universal kit. Integrity of the isolated RNA was analyzed using the Agilent TS HS RNA Kit and TapeStation 4200 at the Genome Technology Center at the Azrieli Faculty of Medicine, Bar-Ilan University. 1000ng of total RNA was treated with the NEBNext poly (A) mRNA Magnetic Isolation Module (NEB, #E7490L). RNA sequencing libraries were produced by using the NEBNext Ultra II RNA Library Prep Kit for Illumina (NEB #E7770L). Quantification of the library was performed using a dsDNA HS Assay Kit and Qubit (Molecular Probes, Life Technologies) and qualification was done using the Agilent TS D1000 kit and TapeStation 4200. 250nM of each library was pooled together and diluted to 4nM according to the NextSeq manufacturer’s instructions. 1.6pM was loaded onto the Flow Cell with 1% PhiX library control. Libraries were sequenced with the Illumina NextSeq 550 platform with single-end reads of 75 cycles according to the manufacturer’s instructions. Sequencing data was aligned and normalized (reads per million mapped reads) using Partek bioinformatics software. Pathway analysis was performed using the ShinyGO webtool(*26*). Heat maps, principal component analysis (PCA) plots and volcano plots were generated using GraphPad Prism software.

### Histology, immunohistochemistry, and inflammation scoring

Distal colon tissues were fixed in 4% paraformaldehyde, paraffin embedded, sectioned, and stained with hematoxylin and eosin. Histopathological scoring, including inflammation and ulceration scores, were performed in a blind fashion by an expert in toxicologic pathology. Immunohistochemistry for neutrophil elastase and myeloperoxidase were preformed using Bioss bs-6982R and Abcam ab9535 antibodies, respectively, as previously described(*27*).

### Blood cytometry and chemistry

Whole blood was drawn via cardiac puncture. Blood samples were collected into a blood collection tube, 50ul for complete blood count (CBC) and 180ul for biochemistry. The samples sent at a temperature of 4 degrees to a service and analysis laboratory: American Medical Laboratories (AML) in Herzliya, Israel.

### Microbiota profiling and analysis

Bacterial DNA was extracted from feces, using the Mobio PowerSoil DNA extraction kit (MoBio) following a 2-minute bead-beating step (Biospec). The V4 of the 16S rRNA gene was amplified using PCR with barcoded primers. DNA was then purified using AMPURE XP magnetic beads (Beckman Coulter) and quantified using Quant-iT PicoGreen dsDNA Assay (ThermoFisher) and equal amounts of DNA were then pooled and sequenced. After sequencing on an Illumina MiSeq platform at the Faculty of Medicine Genomic Center (Bar Ilan University, Safed, Israel), single end sequences reads were import and demultiplexed using QIIME 2 (version 2023.2)(*28*). Sequencing errors were corrected by DADA2(*29*) and taxonomic classification was done using Greengenes reference database with confidence threshold of 99%(*30*). Principal Coordinate Analysis (PCoA) was performed using Jaccard distances, which calculate the difference between the presence or absence of features(*31*). Analysis of composition of microbiome (ANCOM) was used to identify differentially abundant taxa(*32*).

### qRT-PCR

Bacterial DNA was extracted from feces using the Mobio PowerSoil DNA extraction kit (MoBio) following a 2-minute bead-beating step (Biospec). The DNA quantified using Quant-iT PicoGreen dsDNA Assay (ThermoFisher). The concentrations of the samples were diluted to 100ng. qPCR was performed with Cyber Green with the following primers:

#### Pan Bacterial 16s

Forward: 5’-CCTACGGGAGGCAGCAG-3’,

Reverse: 5’-ATTACCGCGGCTGCTGG-3’.

#### Clostridia

Forward: 5’- ACTCCTACGGGAGGCAGC-3-3’,

Reverse: 5’- GCTTCTTTAGTCAGGTACCGTCAT -3’.

### Detection of hypoxia *in vivo*

Mice were treated intraperitoneally with a 1.2mg solution of pimonidazole HCl, (Hypoxyprobe Kit) 1 hour before euthanasia. Distal colon tissues were fixed in 4% paraformaldehyde, paraffin embedded, sectioned, and stained with of 11.23.22.r Rat Mab for 1 hour. Sections were visualized with a Zeiss Axio imager M2 according to manufacturer instructions. Scoring was performed in a blinded fashion.

### Carboxypeptidase -inhibitor treatment

Mice were infected orally with *Salmonella* as above. After infection, mice were treated with 1.25mg of DL-Benzylsuccinic acid (in 10% ethanol) via intraperitoneal injection every 8h every 12h. Control mice were treated with 10% ethanol in PBS. Mice were physically examined before and after injections to exclude damage due to the procedure. Mice hurt during the injection were excluded from the experiment.

### Statistical analysis

Results were analyzed using Graph-Pad Prism 8 software (GraphPad, Inc., La Jolla, CA, USA).

## Supplementary Figures

**Figure S1:**
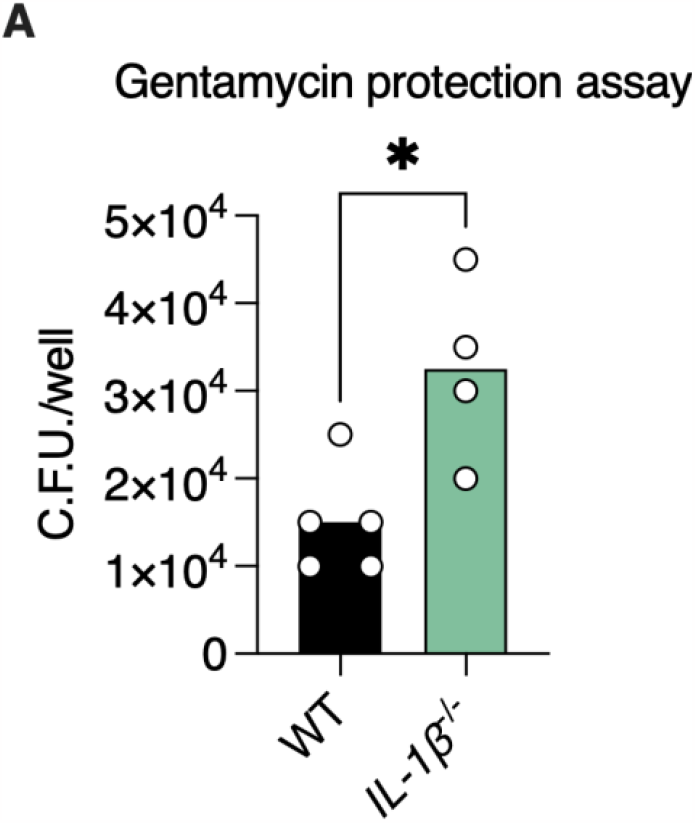
Macrophages from *IL-1β* ^*-/-*^ mice are compromised in their ability to clear intracellular *Salmonella*. (**A**) Gentamycin protection assay using peritoneal macrophages extracted from mice infected with *Salmonella ex vivo* with a MOI of 3. \**P*<0.05; Student’s t-test.

**Figure S2:**
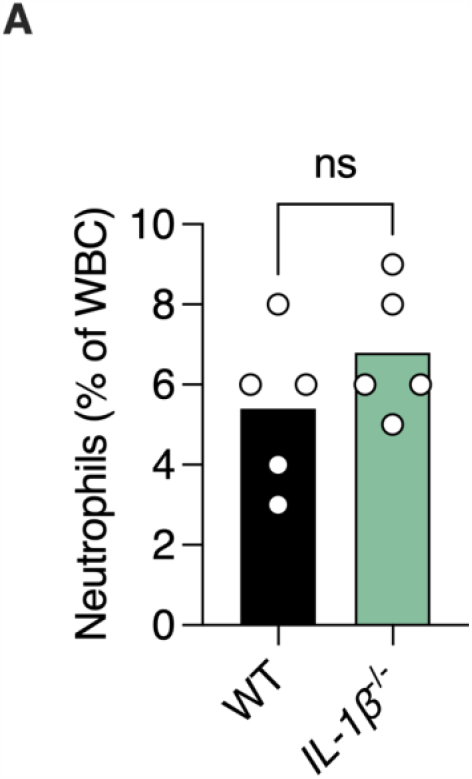
Loss of IL-1β does not affect levels of circulating neutrophils in naïve mice. (**A**) % of neutrophils out of total circulating WBC. ns, not statistically significant; Student’s t-test.

**Figure S3:**
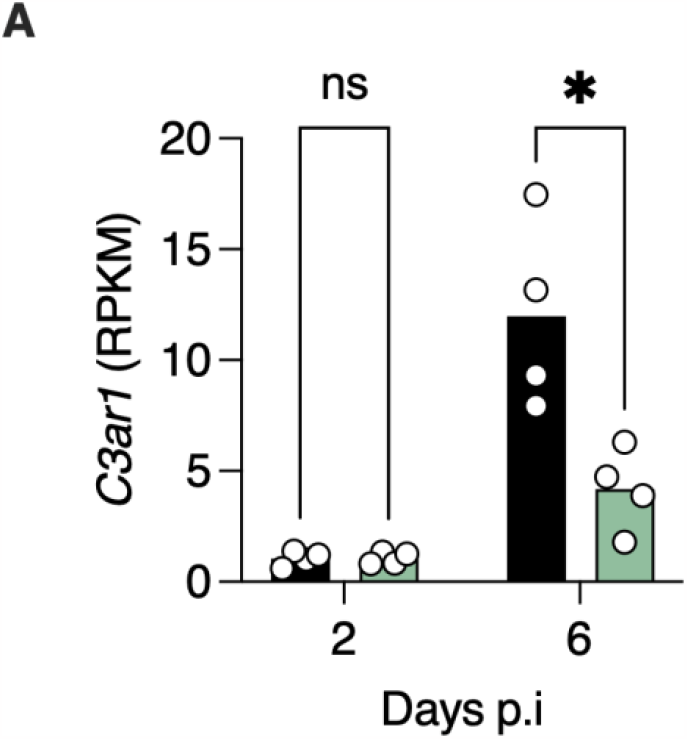
Elevated levels of Complement 3-receptor in *Salmonella*-infected WT mice. (**A**) Normalized reads of the indicated genes from colons of mice based on RNA sequencing. Each dot represents a mouse. \**P*<0.05, ns, not statistically significant; Student’s t-test.

**Figure S4:**
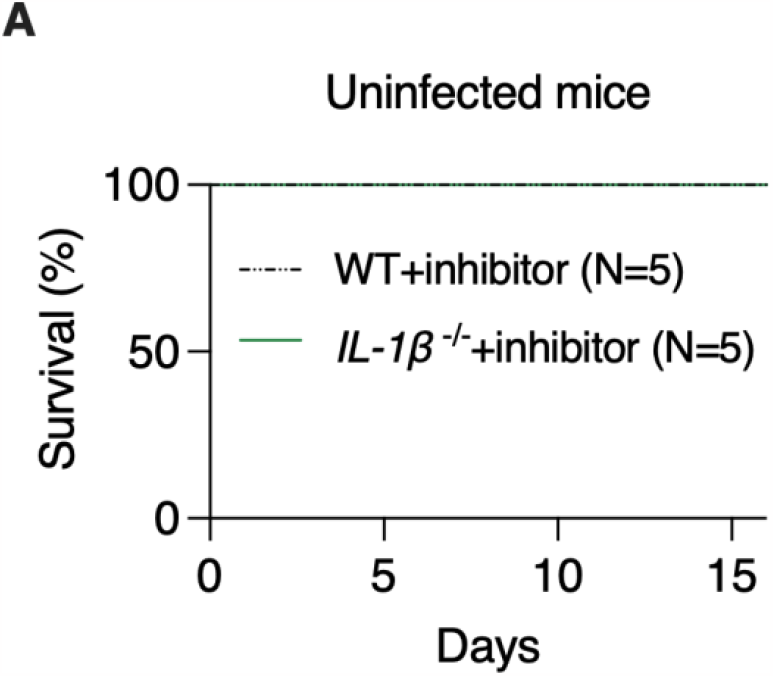
Treatment with carboxypeptidase inhibitor is not lethal in mice. (**A**) Survival of mice treated with carboxypeptidase inhibitor.

